# Dynamic shifts in brain criticality support cognitive processing

**DOI:** 10.64898/2026.03.12.711394

**Authors:** Hongyu Chang, Weiwei Chen, Lindsay A Karaba, Xiyu Mei, Ryan E Harvey, Wenbo Tang, Antonio Fernandez-Ruiz, Azahara Oliva

## Abstract

Systems operating near their critical point, or close to a transition between order and disorder, have computational advantages. In the case of neural networks, proximity to criticality is proposed to support optimal brain function. However, different cognitive processes rely on disparate computational demands. Using large-scale electrophysiological recordings in behaving rodents, we examined how critical dynamics in the hippocampus are regulated during learning and sleep-dependent memory consolidation. We found that operating near criticality enables learning by facilitating hippocampal coordination with input regions and maximizing flexibility of neural representations. In contrast, the hippocampal network shifts toward a more ordered, subcritical regime during sleep memory replay, and recovers its proximity to criticality through cholecystokinin interneurons-mediated inhibition. Overall, our findings provide a biophysical substrate for understanding how critical dynamics in neuronal networks can support a variety of brain functions. Importantly, our results suggest that optimal learning systems, whether biological or artificial, may require a dynamic regulation between flexible and rigid states, and can offer biophysical constraints to guide the design of Large Language Models (LLM) tuned to criticality.

---

Systems closer to their critical point, that is, near a transition between order and disorder, are endowed with several advantages, including a higher dynamic range ^1–3^, enhanced computational capacity ^4–7^, and greater flexibility for information processing ^8–11^. Thus, an influential proposal suggests that the brain operates close to criticality to achieve optimal performance ^6,7,9,12–19^. Physiological signatures of critical dynamics, such as power law distributions, have been observed through *in vitro* preparations ^2,8,20^ and in coarse-grained neural recordings in animals ^21–28^ and humans ^29–36^. Furthermore, several brain disorders are associated with a disruption in critical dynamics ^14,17,18,37–41^. However, it is not yet known what the biophysical nature of criticality in the brain is. Furthermore, different cognitive processes require distinct computational demands. For learning new information, neuronal networks may benefit from higher flexibility or an ample dynamic range, both facilitated near the critical point. In contrast, consolidating recent experiences may require minimizing susceptibility to external perturbations, conditions which will be favored far from the critical point. In this study, we used large-scale electrophysiological recordings in behaving rodents to investigate the neural mechanisms that regulate critical dynamics in the CA1 region of the hippocampus during learning and sleep-dependent memory consolidation. We hypothesized that the hippocampal network adjusts dynamically across distinct critical regimes to support various cognitive processes. Because the critical point is thought to provide systems with a higher dynamic range ^1–3^, optimized computational capacity ^4–7^, and greater flexibility for information processing ^8–11^, we reason that the hippocampus would operate near criticality to optimize learning. Conversely, transitioning into a more subcritical regime during memory consolidation would favor a more ordered state that could facilitate memory replay (Fig. 1A).

**Figure 1.**
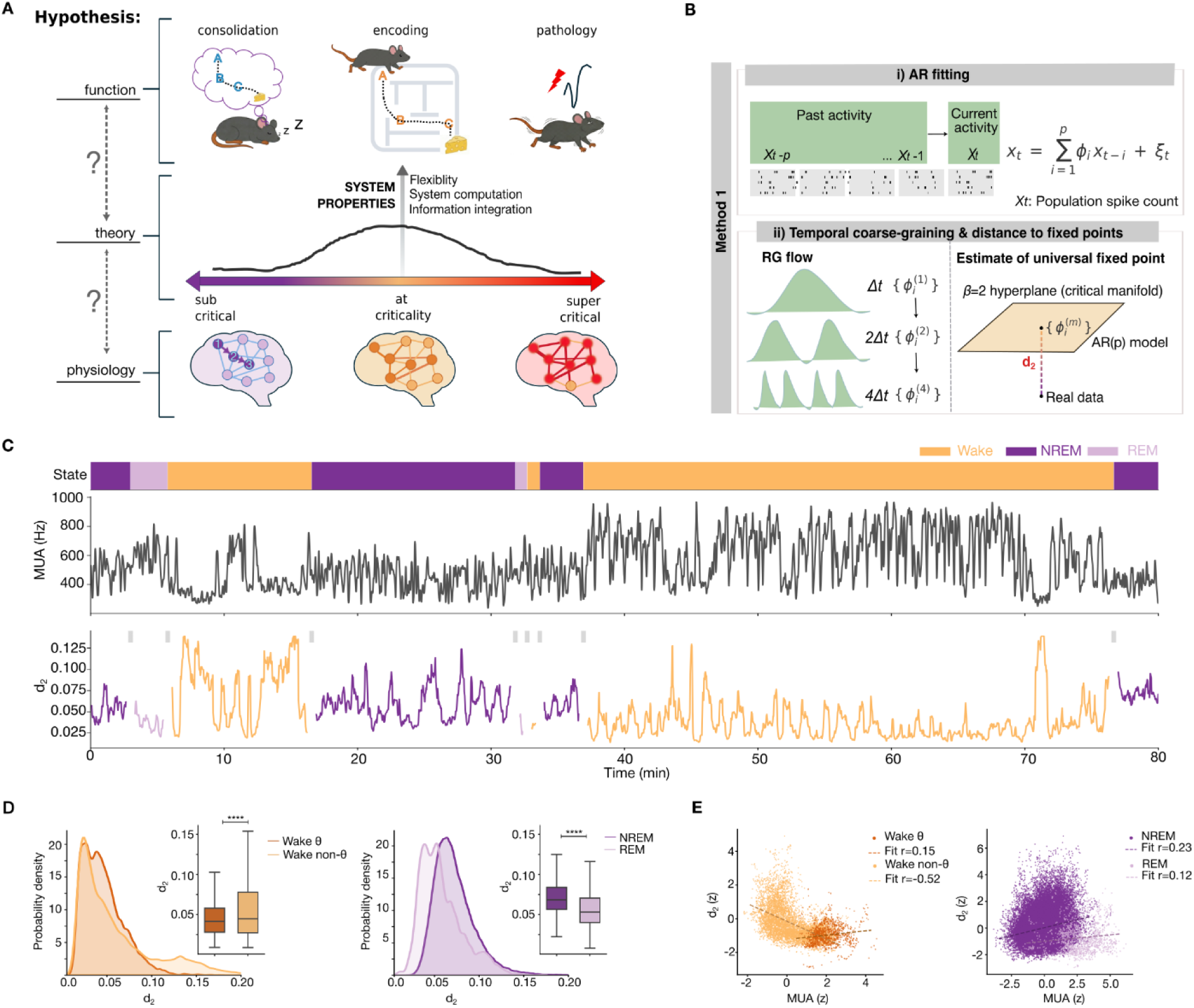
(**A)**. Hypothesis: critical dynamics in the hippocampus fluctuate between a state close to criticality during memory encoding to a far from criticality, subcritical regime during memory consolidation. **(B).** Overview of the methodological approach used to quantify distance to criticality. Top: autoregressive models, AR(p), are fit to the z-scored multiunit spike counts, using the temporal renormalization group (tRG) theory to extract the AR coefficients that are later used to provide a time-resolved estimate of the distance to criticality as d_2_ (bottom), that is, the Euclidean distance from the fitted AR model to the β = 2 fixed-point hyperplane (considered the scale-invariant critical manifold). Note that lower and higher d_2_ values imply closer and farther away from criticality respectively. Grey rectangles denote periods around state transitions during which d_2_ was not computed. **(C).** Example of continuous recording illustrating state segmentation (Wake, NREM, REM; top), multiunit firing rate (middle), and d_2_ to criticality (bottom). **(D).** Left: distribution of d_2_ during waking for active (theta, θ) and non-active (non-theta, non-θ) periods. Inset boxplots show group comparisons of d_2_ value (p < 1×10⁻¹⁵, paired linear mixed-effect model (LMM), n = 18 sessions from n = 6 animals). Right: same for sleeping periods of NREM and REM states. Inset boxplots show group comparisons (p < 1×10⁻¹⁵, paired LMM, n = 18 sessions from n = 6 animals). **(E).** Relationship between d_2_ and multiunit activity (MUA) during theta and non-theta states during wake (left panel) and REM and NREM stages during sleep (right panel). The d_2_ relationship with MUA displayed a positive (r =0.15, p < 1×10⁻¹⁵) and a negative (r = −0.52, p < 1×10⁻¹⁵) correlation during theta and non-theta wake states respectively, whereas both NREM and REM sleep stages showed a positive relationship (NREM: r = 0.23, p < 1×10⁻¹⁵; REM: r = 0.12, p = 1.17 × 10^−5^). Pearson correlations were used for all comparisons, n = 18 sessions from n= 6 animals). Asterisks denote significance (*p<0.05, **p<0.01, ***p<0.001, ****p<0.0001; n.s., not significant).

## Hippocampal activity distance to criticality varies across brain states

Because memory encoding and consolidation happen during distinct brain states ^42^, we first examined hippocampal CA1 activity across waking and sleep (Fig 1). To quantify criticality, we used a recently developed method based on the temporal renormalization group (tRG) theory ^43^ (Fig. 1B). Traditional approaches based on the detection of avalanches (population bursts surpassing a threshold) such as distance to criticality coefficient (DCC) or branching ratio (BR) (fig. S1A) are computed over long periods of time, compromising the discovery of faster dynamic changes in criticality. An advantage of the tRG-based method is the use of autoregressive (AR) models to fit experimental data, which enables a time-resolved estimate of proximity to criticality (d_2_).

As expected, the explained variance of the AR model fit was higher for shorter time windows and plateaued as window length increased (fig. S1B), yielding comparable signal-to-noise (SNR) trends for 30- to 60-second windows consistent with previous reports ^44^ (fig. S1C). To avoid introducing artifactual fluctuations in d_2_, we excluded periods of state transitions, which involve abrupt shifts in firing dynamics (fig. S1D).

We found that the hippocampus’ distance to criticality fluctuated dynamically during awake and sleep states (Fig. 1C), consistent with recent observations in cortical areas ^25,28,43,45,46^. On average, the hippocampus operated closer to criticality during waking than during sleep (fig. S1E). Specifically, the hippocampus was closer to criticality during active theta (Fig. 1D, left) and during REM (Fig. 1D, right) compared to wake non-theta and NREM sleep respectively. Comparable results were obtained when analyzing the BR, which quantifies how much activity propagates from one time bin to the next (fig. S1F).

Overall, during waking, CA1 firing rates were positively correlated with *d_2_* during theta and negatively correlated during non-theta periods (Fig. 1E, left). In contrast, both REM and NREM sleep showed positive correlations between firing rates and *d_2_* (Fig. 1E, right). Because one of the hallmarks of criticality is a higher dynamic range of the system ^1–3^, we next quantified the dynamic range of CA1 firing rates ^2^ in relation to d_2_. Notably, despite state-dependent differences in d_2_, we found a higher dynamic range of CA1 firing rates during both awake and sleep states (fig. S1G, left and right respectively) when it was closer to criticality. Overall, these results indicate that hippocampus’ critical dynamics fluctuate rapidly within each brain state and strongly differ across states, while maintaining a consistently higher dynamic range in proximity to criticality.

## The hippocampus moves away from criticality during replay of recent experiences

Because the hippocampus moved away from criticality during sleep, we sought to understand whether this departure was related to sleep-dependent memory consolidation processes, as we originally hypothesized (Fig. 1A). To answer this, we first compared d_2_ values in sessions of pre-versus post-task sleep in mice that trained on a hippocampal-dependent spatial memory task ^47^. We found that d_2_ distance increased during post- compared to pre-task NREM sleep (Fig. 2A). This pre- to post-task NREM sleep increase was more pronounced after animals trained in a novel maze than following training on a familiar maze (fig. S2A), suggesting that higher memory load displaced the network further from criticality.

**Figure 2.**
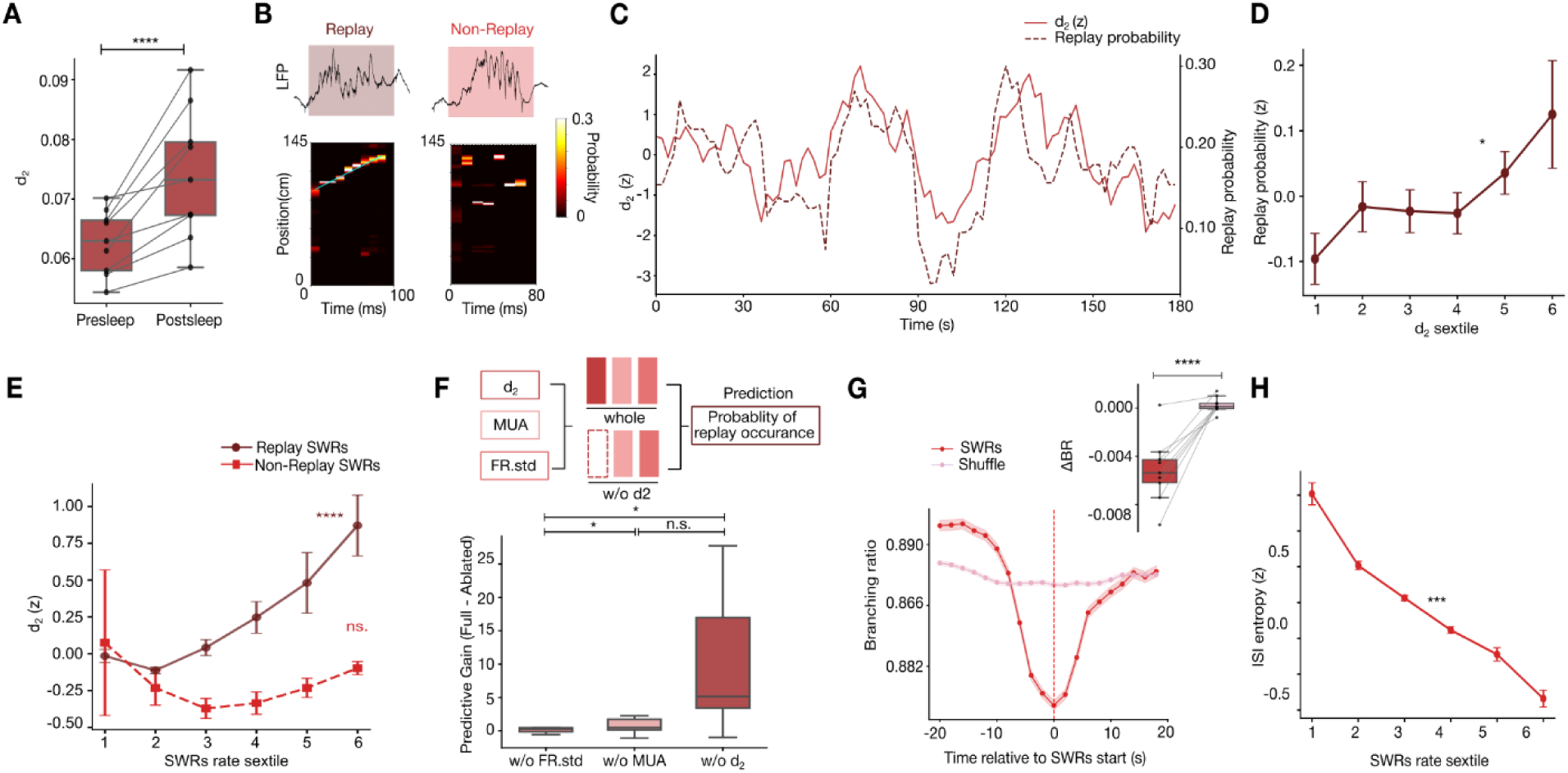
The hippocampus moves away from criticality during memory replay. **(A).** d_2_ increases from pre- to post-task sleep (p = 2.36 × 10⁻⁶, n = 9 sessions from n = 6 animals, paired LMM). **(B).** Example of SWRs associated with replay (ordered firing, left) and non-replay (no preferred order, right). For each event CA1 LFP and decoded position (the cyan line denotes the best linear fit) for the event are shown **(C).** Example trace of z-scored d2 (red, left Y axis) and replay probability (dashed dark red, right Y axis), computed in 30-s sliding windows. **(D)**. Z-scored replay probability (mean ± SEM) per d_2_ sextiles of NREM sleep. Note that replay probability increases with higher d_2_ values (p = 0.016, n = 9 sessions from n = 6 animals; linear mixed-effect model). (**E).** d_2_ correlated with the rate of SWRs associated with replay but not with the rate of SWRs not associated with memory replay (p < 10^−15^ and p = 0.80 for replay and non-replay curves respectively, n = 9 sessions from n = 6 animals, linear mixed model). (**F).** Top: schematic of the logistic regression framework used to predict replay occurrence from predictors (d_2_, MUA, FR_std) while allowing variability of feature effects across bins as a feature×state interaction term. Bottom: predictive gain for each feature interaction term, quantified by the change in cross-validated performance upon feature ablation. Post hoc paired Wilcoxon tests (Holm-corrected) indicated d_2_ > MUA (p = 0.035) and d_2_ > FR_std (p = 0.039), while MUA ≈ FR_std (p = 0.91). **(G)**. Branching ratio (BR) during SWRs and shuffled periods for an example session. Ripple windows showed a significant decrease in BR compared to shuffles (p < 1×10⁻¹⁵, n = 9 sessions from n = 6 animals, paired LMM). (**H).** Inter-spike-interval (ISI) entropy decreased with ripple rate (Pearson’s: r = −0.99, p = 3.57×10^−4^, n = 9 sessions from n = 6 animals).

A well-established mechanism for sleep-dependent memory consolidation is the replay of neural sequences corresponding to activity patterns expressed during recent behavior ^47–53^. Such memory replay is orchestrated by hippocampal sharp-wave ripples (SWRs) ^49,54^, whose incidence increases after learning (fig. S2B) ^53,55^. Therefore, we detected replay events (Fig. 2B) during sleep using a Bayesian decoder trained on the CA1 activity patterns recorded during the behavioral task ^47,56^. Approximately 22 % of SWRs were associated with significant replay (Fig. 2B), consistent with previous reports ^47,52,56–59^.

Interestingly, we observed that replay probability co-fluctuated with the hippocampus’ distance to criticality during NREM sleep (Fig. 2C), and that the two measures were significantly correlated (Fig. 2D). Conversely, d_2_ was positively correlated with the rate of replay events but not with the rate of SWRs lacking significant replay (Fig. 2E), and this dissociation couldn’t be explained by the associated d_2_ uncertainty ^44^ within SWR’s sextiles (fig. S2C).

To quantify the relative contribution of different variables to replay probability, we built a generalized linear model (GLM) (Fig. 2F, top). The full model *(whole*) was fitted with a combination of distance to criticality (*d_2_*), mean multiunit firing rate (*FRm*) and multiunit firing rate variability across neurons (*FRstd*) during NREM sleep. Ablation models were generated by removing different variables. The reduction in predictive gain was significantly greater when *d_2_* was ablated compared to *FRm* or *FRstd* ablation (Fig. 2F, bottom), suggesting that proximity to criticality is a strong predictor of replay probability.

As an alternative approach of assessing criticality, we quantified avalanche size and duration (fig. S2D), power-law scaling properties (fig. S2E) and DCC (fig. S2F) ^25,45^. For all three measures, we found that NREM periods of high-SWR rate, during which replays were most prominent ^47^, were farther from criticality than periods of low-SWR rate (fig. S2D, E, F).

Next, we asked whether the network shifted towards sub- or super-critical regimes around SWRs. To answer that, we calculated the BR (fig. S1A), which quantifies how activity propagates from one time bin to the next and classifies dynamics as subcritical (BR < 1), critical (BR ≈ 1), or supercritical (BR > 1) ^8,24,60^. The BR in CA1 showed subcritical values and significantly decreased during periods of high SWR rate compared to shuffles (Fig. 2G). Because subcritical regimes are characterized by lower entropy compared to critical regimes ^9,11^, we next quantified the entropy of CA1 firing dynamics across different SWR rates. We found that entropy was significantly lower during periods of higher SWR rate (Fig. 2H).

Our observation that the hippocampus moves away from criticality during post-task NREM sleep raises two possible interpretations. One is that the structured population spiking bursts associated with memory replay drives the observed elevated d_2_ values. Alternatively, the observed increase of d_2_ values could reflect a global transition into a subcritical regime, which may itself be a prerequisite for replay. To distinguish between these possibilities, we first checked for the increased SWR rate after learning (fig. S2B) by comparing pre- and post-task NREM periods of matched SWR rate distributions. We found that d_2_ remained higher in post- compared to pre-task NREM sleep after controlling for SWR rate (fig. S3A). This result persisted even after excluding all spikes within SWRs and analyzing only the remaining activity within the time window (fig. S3B). Similarly, the positive correlation between SWR rate and d_2_ remained after removing all spikes within SWRs (fig. S3C). Similarly, the entropy of the firing dynamics continued to show a negative correlation with SWR rate after removing SWR firing (fig. S3D). Altogether these results show that during post-task NREM sleep, CA1 transitions into a highly subcritical regime that provides a permissive network state for the emergence of memory replay during SWRs in support of memory consolidation.

## BARRs contribute to restoring the hippocampal network to its critical point

Because we observed that the hippocampus moved away from criticality during memory consolidation (Fig. 2), we next asked how the hippocampal network restores its proximity to criticality. To identify potential mechanisms able to restore proximity to criticality, we built a simplified CA1 network model. To investigate how the activity of distinct neuronal populations influences critical dynamics beyond excitation and inhibition, the model included pyramidal cells (E), fast-spiking parvalbumin (PV) interneurons, and regular-spiking cholecystokinin-expressing (CCK) interneurons ^61–65^. In addition, it featured excitatory inputs from CA3, known to be important for SWRs generation, targeting all cell types in CA1 ^64^ and from CA2 targeting CA1 CCK interneurons ^53^ (Fig. 3A).

**Figure 3.**
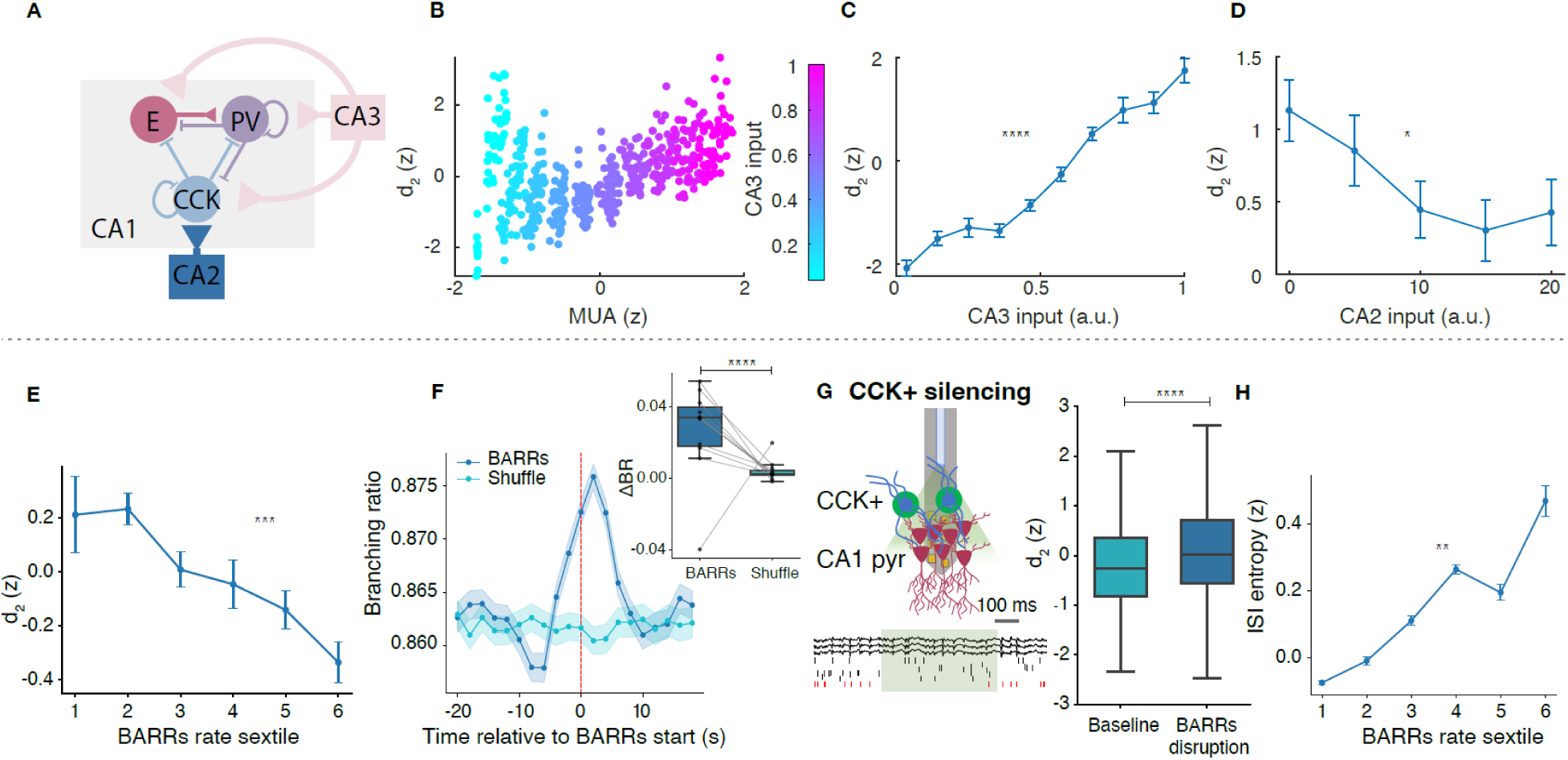
CCK inhibition during BARRs contributes to restoring criticality in the hippocampus. **(A)**. Simplified network model of CA1, featuring excitatory (lines with triangles) and inhibitory connections (T-lines), and neuronal subpopulations in different colors. The model comprised one recurrent layer with a pyramidal neurons’ population (E) and two interneuron populations (PV and CCK), as well as two feedforward inputs from CA3 and CA2, modeled as independent Poisson units, providing excitatory input to the CA1 network. (**B).** In the model, CA1 d_2_ was significantly correlated with multiunit firing rate (Spearman ‘s: ρ = 0.92, p < 10^−15^), which was in turn modulated by the strength of CA3 input. (**C).** CA1 d_2_ increased with the strength of CA3 input (p < 10^−15^, r = 0.92, Spearman correlation). **(D)**. CA1 d_2_ decreased with the strength of CA2 input (p = 0.011, r = −0.35, CA3 Input = 0.9, Spearman correlation). **(E).** In mice, CA1 d_2_ was negatively correlated with the rate of barrages (BARRs) (Pearson’s r = −0.969, p = 0.0014 mean ± SEM). **(F).** Branching ratio (BR) during BARRs and shuffled periods for an example session. BARR windows showed a significant increase in BR compared to shuffles (p = 2.71×10⁻^7^, n=11 sessions from n = 5 animals, paired LMM). **(G).** Left: schematic of optogenetic inhibition of CA1 CCK basket cells during sleep. Right: d2 stayed larger in sessions with CCK inhibition compared to control sessions (p < 10^−15^, n=7 session from n = 3 animals, paired linear mixed model). **(H).** Inter-spike-interval (ISI) entropy was positively correlated with the rate of BARRs (Pearson’s: r = 0.946, p = 0.0043, n = 12 session from n = 5 animal, mean ± SEM).

The model reproduced the relationship between multiunit firing rate and d_2_ experimentally observed during sleep (fig. S4, Fig. 3B and Fig.1 E, right). The strength of CA3 input was positively correlated with d_2_ in CA1 (Fig. 3C), consistent with our experimental results during SWRs (Fig. 2D, E and fig. S3F). In contrast, the strength of CA2 input onto CCK interneurons was negatively correlated with d_2_ (Fig. 3D). This result led us to hypothesize that CCK-mediated inhibition in CA1 may provide a mechanism for restoring the network’s proximity to criticality.

Recently, we showed that a circuit from CA2 pyramidal cells to CCK basket cells in CA1 generates barrages of action potentials (BARRs) during sleep to counterbalance the elevated firing rates associated with memory replay (fig. S4A) ^53^. Therefore, we next asked whether BARRs contribute to restoring the network’s proximity to criticality. To answer that, we performed simultaneous recordings from CA1 and CA2 regions during the same behavioral paradigm and following the same task structure than before (Fig. 3E-H). We identified BARR events during post-task sleep, which, as expected, were accompanied by increased CA2 pyramidal cell activity (fig. S3G). BARR rate was inversely correlated with d_2_ during post-task NREM sleep (Fig. 3E), in line with the predictions from our model (fig. S4B-D, E-G, H-J). Similarly, the BR significantly increased during BARRs compared to shuffles (Fig. 3F), even when removing all spikes within BARRs (fig. S3E), and the entropy of the CA1 firing dynamics was larger during periods of higher BARRs’ rate (Fig. 3H).

To causally test the contribution of CCK-mediated inhibition to network dynamics, we next looked at sessions during which a subset of CA1 CCK interneurons was optogenetically silenced during BARRs in post-task sleep ^53^. In these sessions, the CA1 network remained farther away from its critical point compared to control sessions (Fig. 3G). Together, these results demonstrate that CCK-mediated inhibition during BARR states contribute to restoring the hippocampal network’s proximity to criticality.

## Proximity to criticality of hippocampal activity facilitates learning

Next, we sought to test our prediction that proximity to criticality facilitates learning (Fig. 1A). To do that, we turned to a spatial task that enabled us to assess de novo learning each day. We performed silicon probe recordings of all subregions of the hippocampus and medial entorhinal cortex (MEC) in rats during a spatial learning task in a cheeseboard maze. In this task, rats had to learn the location of three hidden water rewards whose configuration changed daily ^59,66–68^, followed by ∼1.5 hours of sleep in their home-cage (Fig. 4A). A delay period in the start-box was imposed between trials during learning. On the following day, animals underwent a recall test to evaluate memory for the previous day’s reward locations ^59,66–68^ (Fig. 4A).

**Figure 4.**
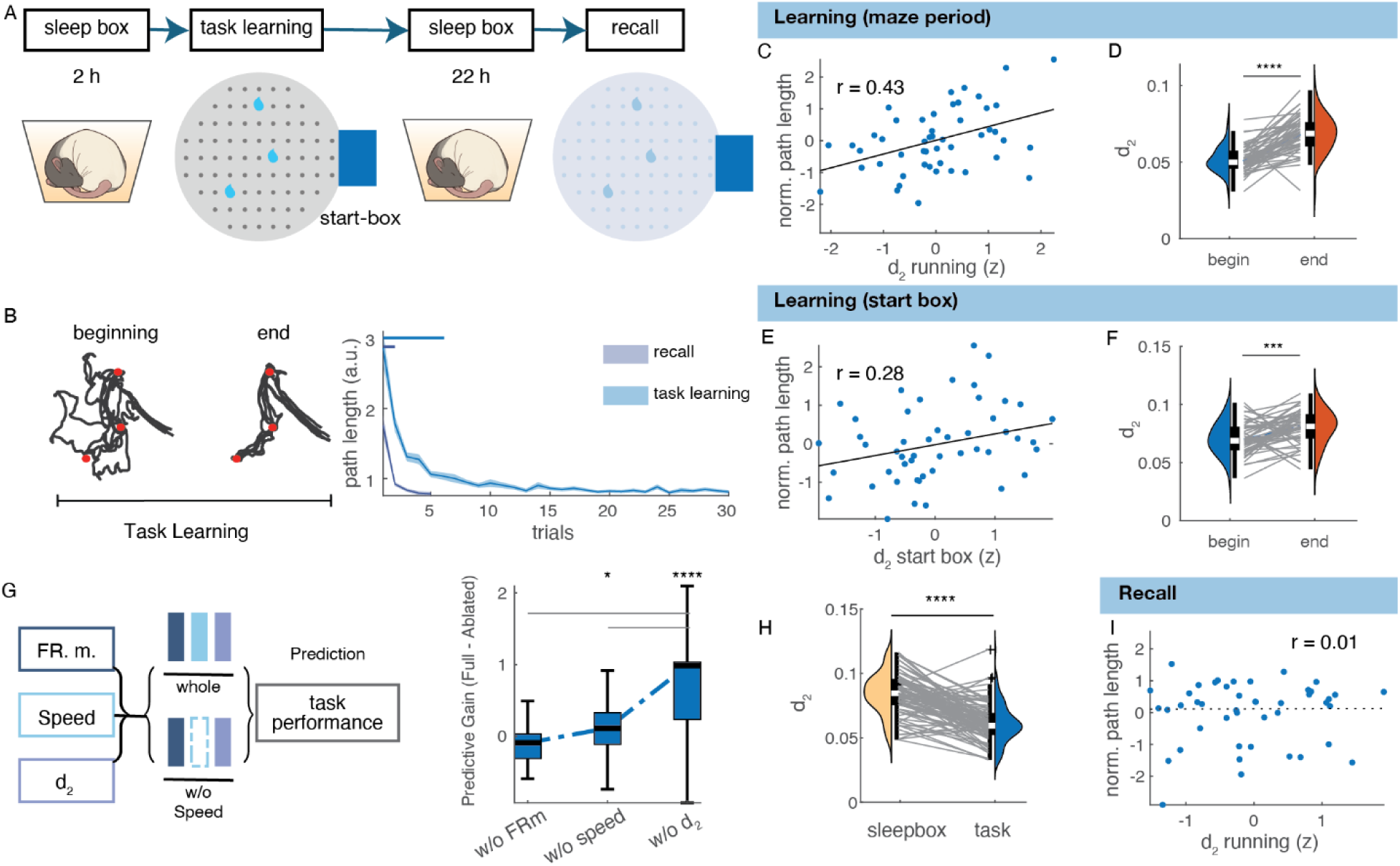
Hippocampus proximity to criticality predicts behavioral performance. **(A)**. Schematic of the cheeseboard task. Rewards’ location changed daily, forcing the animals to learn a new path each day, and memory consolidation was tested 22 hour later. **(B)**. Left: example trajectories in early and late trials. Red dots = reward locations. Behavioral performance (assessed as mean path length) during the task learning and recall phases. Animals learned the optimal trajectory to collect all three rewards within the first few trials of the task (p < 10^−15^, n = 49 sessions from n = 3 animals, Kruskal-Wallis test with post-hoc Dunn test and Sidák’s adjustment against last trial) and first 2 trials during the recall phase (p < 10^−15^, n = 45, Kruskal-Wallis test with post-hoc Dunn test and Sidák’s adjustment against last trial). **(C)**. Behavioral performance was correlated with d_2_ during the earlier trials (first 5 trials) of the learning phase (r = 0.43; Spearman: r = 0.39, p = 5.90×10^−3^). **(D)**. d_2_ was lower in the earlier (first 5 trials) compared to later (last 5 trials) trials during the learning phase of the task (p=1.50×10^−8^, n = 49, Wilcoxon signed rank test). **(E)**. Behavioral performance was correlated with d_2_ during the preceding start-box periods (r = 0.28; Spearman: r = 0.32, p = 0.026). **(F)**. d_2_ during start-box periods was significantly lower in the earlier compared to later trials (p = 9.46×10^−4^, n = 49, Wilcoxon signed rank test). **(G)**. Left: Schematic of the GLM model. Right: prediction gain of the full model compared with different ablation models. Note the strongest reduction of predictive gain for d2 (*p=0.023, ****p= 1.316×10^−13^, Wilcoxon signed rank test). **(H)**. d_2_ during the task was lower compared to sleep-box waking periods of equal duration (5 minutes periods; p < 0.0001, Wilcoxon signed rank test, n = 90 sessions from n = 4 animals). **(I)**. Behavioral performance was not correlated with d_2_ during the recall phase (r = 0.01; Spearman: r = 0.3, p = 0.84).

We replicated our previous findings in this new dataset, including that CA1 operated closer to criticality during theta compared to non-theta waking periods, and during REM relative to NREM sleep (fig. S5A, B). The dynamic range of CA1 firing rates was also higher when the hippocampus was closer to criticality (fig. S5C, D). In addition, we observed that the hippocampus departed and returned closer to criticality around SWRs (fig. S6A-D) and BARRs (fig. S6E-G) respectively, paralleled by a decrease and increase of the neuronal inter-spike-interval entropy.

During the task, rats engaged in random search during the earlier trials and quickly learned to take directed trajectories towards rewarded locations (Fig. 4B). We hypothesized that a network regime closer to criticality would benefit subsequent learning on this task. To test this, we compared behavioral performance (quantified as path length to collect all rewards) with CA1 distance to criticality measured across different task phases.

We found that proximity to criticality during learning trials was positively correlated with performance (Fig. 4C and fig. S7). Overall, the hippocampus was closer to criticality during earlier compared to later trials (Fig. 4D), and these effects could not be explained by speed, firing rate or theta power (fig. S7). Moreover, the hippocampus’ distance to criticality during inter-trial intervals in the start-box correlated with subsequent trial performance (Fig. 4E), and inter-trial intervals during earlier trials were closer to criticality compared to later trials (Fig. 4F).

To assess the relative contribution of different variables to learning performance, we constructed a generalized linear model (GLM) (Fig. 4G, left). The full model *(whole*) incorporated multiunit firing rate (*FRm*), animal speed (*speed*) and distance to criticality (*d_2_*) during the task. We then generated ablation models by removing each one of the predictors. We found that the largest drop in predictive gain between the whole and the ablated models was obtained for *d_2_* (Fig. 4G, right), indicating that proximity to criticality strongly contributes to predicting behavioral performance. Finally, we asked whether these results were specific to learning or instead reflected a general property of the waking state. To address this, we compared home-cage waking periods with inter-trial start-box intervals. The CA1 network was consistently closer to criticality during start-box periods than during home-cage waking epochs of the same duration (Fig. 4H). In addition, proximity to criticality no longer predicted performance during next day recall trials (Fig. 4I) after animals had already learned the reward configuration, in contrast to the learning phase (Fig. 4C, D).

To investigate the neural circuit mechanisms linking criticality in the hippocampus with behavioral performance, we examined how strongly CA1 coordinated with upstream inputs. To quantify cross-regional coordination between different areas, we computed canonical correlation analysis (CCA) between CA1 and CA2, CA3, or MEC regions (fig. S8A) during the earlier learning trials (Fig. 5b). CCA between MEC and CA1 showed a negative correlation with distance to criticality, indicating a higher coordination between MEC and CA1 when the CA1 network was closer to criticality. A similar result was found between CA1 and CA3, as well as between CA1 and CA2 (fig. S8B, C-E). Overall, these results show that CA1 network can more effectively coordinate with input regions when operating closer to criticality.

**Figure 5.**
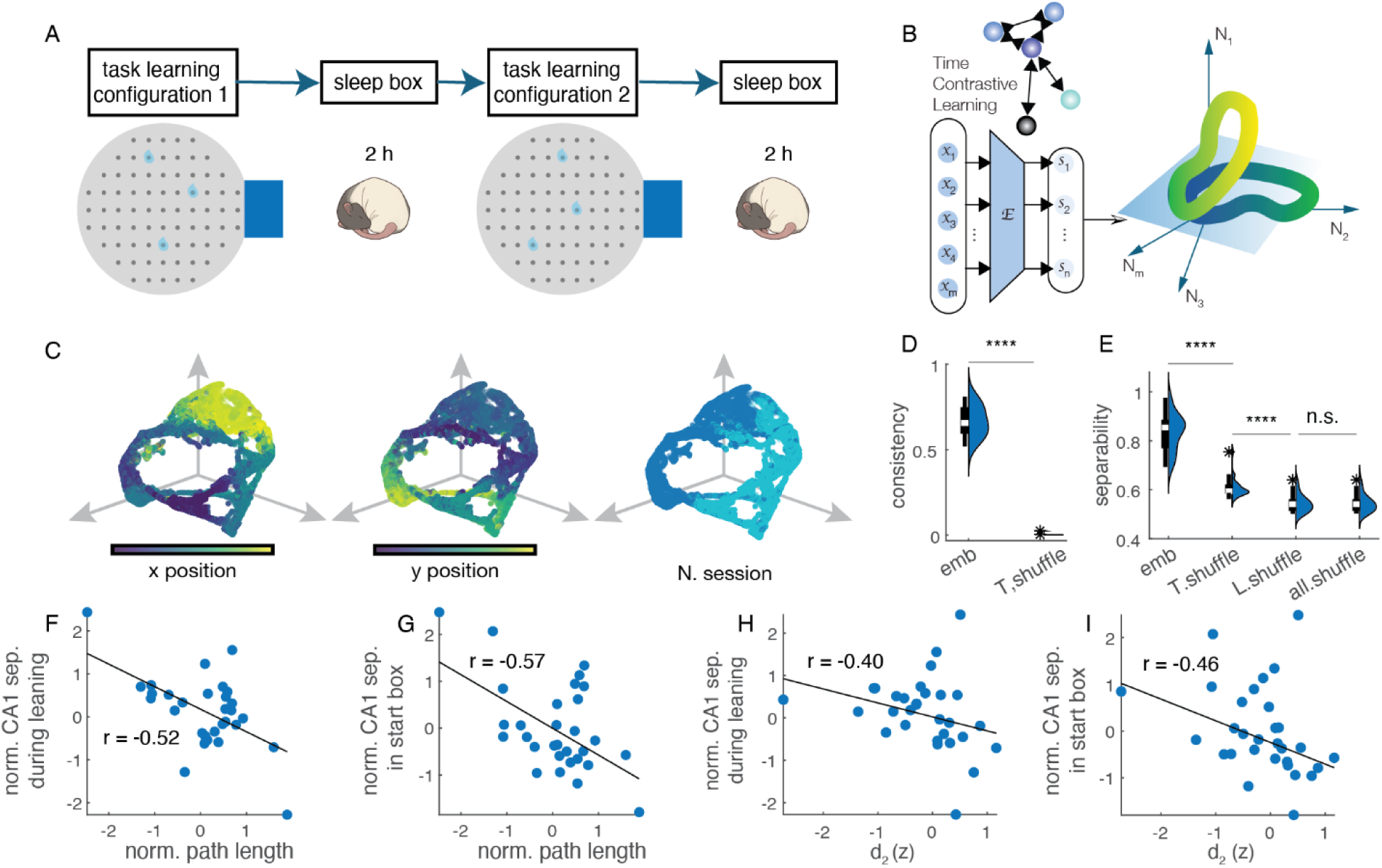
Flexibility of hippocampal representations emerges near criticality. **(A)**. Schematic of the two-configurations cheeseboard task. **(B)**. A time contrastive learning-based method was used to find the latent hippocampal representations of the different task configurations. **(C)** Left and middle: color coded manifold for X and Y positions in the cheeseboard maze. Right: Light and dark blue colors representing the two maze configurations. **(D)** Latent embedding consistency is significantly higher than time shuffled embedding. latent manifold consistency measured as R^2^ of the linear fit between 20 times of randomly trial shuffled embeddings (p = 2.70×10^−5^, n = 30, Wilcoxon signed-rank test). **(E)**. A support vector machine (SVM) was trained to separate manifolds representing the two configurations. Separability was measured as accuracy for each session with 5-fold cross-validation and compared to separability based on time sample shuffled embeddings, task label shuffled data, or time and label shuffled data (p = 1.82×10^−5^, n.s. : p = 0.63, Wilcoxon signed-rank test, n = 30 sessions). **(F)** Separability of neural representations in CA1 during learning phase of the second configuration was correlated with behavioral performance (r = −0.52; Pearson: r = −0.58, p = 8.54×10^−4^, n = 30 sessions). **(G)** Separability of neural representations in CA1 during start-box periods in the second configuration was correlated with behavioral performance (r = −0.57; Pearson: r = −0.54, p = 2.18×10^−3^, n = 30 sessions). **(H)** Separability of CA1 maps during the learning phase of the second configuration was correlated with d_2_ (r = −0.33, Spearman: r = −0.40, p = 0.029, n = 30 sessions). **(I)** Separability of CA1 latent embeddings during start-box periods in the learning phase of the second configuration was correlated with d_2_ (r = −0.46, Spearman: r = −0.40, p = 0.015, n = 30 sessions).

## Distinct hippocampal representations emerge near criticality

Next, we sought to understand the computational mechanism through which proximity to criticality benefits learning. An additional hallmark of systems close to criticality is increased flexibility, that is, the capacity to access a broader range of trajectories in state space in response to inputs ^4,11,69^. Recently, we showed that the hippocampus exhibits a similar form of flexibility, generating distinct latent representations of the same environment upon contextual changes, such as different reward configurations ^68^. To investigate how criticality relates to such representational flexibility, we trained rats in a dual-configuration version of the original cheeseboard maze task ^68^. In this task, rats were trained to collect three hidden rewards located in fixed locations over ∼20-30 trials, followed by a sleep session in the home-cage. They were then introduced to a new reward configuration for another ∼20-30 trials (Fig. 5A). Sleep sessions preceded and followed each training session, and a recall trial was run 20 hours (next day) after the training in the second configuration.

To characterize the degree of hippocampal representational flexibility, we used dimensionality reduction techniques to extract neural representations of the two different reward configurations ^68^. Specifically, we looked at CA1 population activity during learning trials using time contrastive learning implemented with CEBRA (Consistent Embedding for Bipartite Relationship learning with Auxiliary variables) (Fig. 5B), a self-supervised machine learning method to reduce neural activity into a low-dimensional manifold ^70^. Two different maps of the same maze corresponding to the two reward configurations emerged at the population level, consistent with recent findings ^68,71^ (Fig. 5C), while single cell responses showed only moderate changes across maze configurations (fig. S9A). These maps, or neural manifolds, displayed significantly higher consistency than shuffles (Fig. 5d). Similar results were found when assessing manifolds’ separability by training a support vector machine (SVM) to separate the two configurations versus shuffles (Fig. 5E), and the separability quickly saturated when considering low latent dimensions (fig. S9C).

Better behavioral performance in the second session of the task was associated with greater separability of the manifolds of the two task configurations, both during learning trials (Fig. 5F) and during inter-trial intervals (Fig. 5G), and these results couldn’t be explained by a trivial multiunit firing rate relationship (fig. S9D, E).

In addition, the separability of the different maps was larger when the hippocampus operated closer to criticality, whether assessed during learning trials (Fig. 5H) or during inter-trial intervals in the second maze session (Fig. 5I). Importantly, this relationship between proximity to criticality and separability was not present during the first maze session for either running trials or inter-trial intervals in the start-box (fig. S9 H, I). Similarly, proximity to criticality during a recall session preceding the learning during the earlier trials before the first maze did not predict map separability (fig. S9 F, G). Together, these findings demonstrate that greater representational flexibility, quantified as differentiation of neural representations in CA1, emerges in proximity to criticality.

## Discussion

In this study, we investigated how critical dynamics in the hippocampal CA1 network contribute to the circuit mechanisms of memory encoding and consolidation. We hypothesized that a neuronal network close to criticality would facilitate memory encoding during behavior, while subcritical regimes would favor the occurrence of memory replay during sleep (Fig. 1A). First, we found that criticality in the hippocampus fluctuates across and within brain states following a faster temporal dynamic than previously shown ^25,45^. The hippocampus resided closer to criticality during awake compared to sleep (Fig 1), in line with recent results from cortical activity ^43,72,73^. During sleep, we found that the hippocampus moved into its most subcritical regime during memory replay (Fig 2), suggesting that replay generation could be facilitated by a transient ordered state, which can be achieved in a subcritical domain. In support of this, removing all firing during SWRs but preserving the windows around them, did not change CA1 distance to criticality, suggesting that a subcritical state provides a permissive dynamic regime that facilitates replay generation. Using a biophysically realistic model of CA1, we predicted that the proximity to criticality could be restored by an increased activity of CCK interneurons (Fig 3, top). We found that during BARRs, hippocampal network states that entrain a circuit from CA2 to CA1 CCK interneurons ^53^, the hippocampal network relaxes closer to criticality (Fig. 3, bottom). This suggests that BARRs may serve as a homeostatic mechanism for restoring an optimal computational regime, better achieved close to criticality.

Animals’ learning performance correlated with proximity to criticality, and this correlation disappeared once animals had already learned (Fig 4). The contribution of the critical regime to efficient learning could be driven by the enhanced dynamic range of firing rates (fig. S1G), and better coordination with input areas, hallmarks of systems working close to criticality. In support of this, we found a higher coordination of CA1 activity with input regions, including MEC, CA3, and CA2, near criticality (fig. S8). Another proposed advantage of systems that operate near criticality is higher flexibility or access to multiple system states ^4,11,69^. Indeed, we found that representational flexibility, quantified as distinct neural maps associated with different contexts, emerged when the hippocampus operated near criticality (Fig 5). Such neural flexibility in the hippocampus can enable the integration of new memory representations into an existing repertoire ^74^, a mechanism that could facilitate continuous learning ^5,75,76^.

Importantly, although the hippocampus featured distinct critical regimes during memory encoding and consolidation, both states remained below the critical point. In contrast, the hippocampus entered a supercritical regime during epileptic seizures (fig. S10), underscoring the importance of optimal critical dynamics.

In summary, our study establishes a link between the free-scaling properties of a physical system, assessed by critical dynamics, and the biological substrate, assessed by specific circuit mechanisms, that supports brain functions. We showed that memory encoding and consolidation alternate between the vicinity of the critical point and a subcritical regime to balance computational demands and network homeostasis. Our findings suggest that optimal learning systems, whether biological or artificial, may require a dynamic regulation between flexible and rigid states, and can offer biophysical constraints to guide the design of Large Language Models (LLM) tuned to criticality, a current solution used to overcome the problem of information scarcity in artificial intelligence (AI) ^77–79^.

## Acknowledgments

We thank members of the Oliva and Fernandez-Ruiz laboratories for useful comments on the manuscript.

## Author contributions

Conceptualization: A.O., A.F.-R., H.C., W.C.; Investigation: H.C., W.C.; Visualization: H.C., W.C, L.A.K.; Methodology: H.C., W.C, L.A.K., X.M., R.E.H., and W.T.; Funding acquisition: A.O. and A.F.-R.; Supervision: A.O.; Writing – original draft: A.O.; Writing – review and editing: A.O., A.F.-R., H.C., W.C., L.A.K., X.M., R.E.H., and W.T.

## Data and materials availability

All data are available in the main text or the supplementary materials. Custom scripts used in this study can be downloaded from https://github.com/ayalab1/neurocode.

